# Complementary effects of adaptation and gain control on sound encoding in primary auditory cortex

**DOI:** 10.1101/2020.01.14.905000

**Authors:** Jacob Pennington, Stephen David

**Affiliations:** Department of Mathematics, Washington State University Vancouver; Oregon Hearing Research Center, Oregon Health and Science University

## Abstract

An important step toward understanding how the brain represents complex natural sounds is to develop accurate models of auditory coding by single neurons. A common model for auditory coding is the linear-nonlinear spectro-temporal receptive field (LN model). The LN model accounts for many features of auditory tuning, but it cannot account for long-lasting effects of sensory context on sound-evoked activity. Two mechanisms that may support these contextual effects are short-term plasticity (STP) and contrast-dependent gain control (GC), each of which has inspired an expanded version of the LN model. Both of these models improve performance over the LN model, but they have never been compared directly. Thus, it is unclear whether they account for distinct processes or describe the same phenomenon in different ways. To address this question, we recorded activity of neurons in primary auditory cortex of awake ferrets during presentation of natural sounds. We then fit models incorporating one nonlinear mechanism (GC or STP) or both (GC+STP) using this single dataset, and measured the correlation between the models’ predictions and the recorded neural activity. Both the STP and GC models performed significantly better than the LN model, but the GC+STP model performed better than either individual model. We also quantified the similarity between STP and GC model predictions and found only modest equivalence between them. Similar results were observed for a smaller dataset collected in clean and noisy acoustic contexts. These results suggest that the STP and GC models describe distinct, complementary processes in the auditory system.

**Significance Statement:** Computational models are used widely to study neural sensory coding. However, models developed in separate studies are often difficult to compare because of differences in stimuli and experimental preparation. This study develops an approach for making systematic comparisons between models that measures the net benefit of incorporating additional nonlinear elements into models of auditory encoding. This approach was then used to compare two different hypotheses for how sensory context, that is, slow changes in the statistics of the acoustic environment, influences activity in auditory cortex. Both models accounted for complementary aspects of the neural response, indicating that a hybrid model incorporating elements of both models provides the most complete characterization of auditory processing.

## Introduction

The sound-evoked spiking activity of an auditory neuron can be modeled as a function of a time-varying stimulus plus some amount of error, reflecting activity that cannot be explained by the model. Prediction error can reflect experimental noise (physiological noise or recording artifacts), but in many cases it also reflects a failure of the model to account for some aspects of sound-evoked activity (Sahani and Linden 2003). A common encoding model used to study the auditory system is the linear-nonlinear spectro-temporal receptive field (LN) model (David et al. 2007; Calabrese et al. 2011; Rahman et al. 2019). According to the LN model, the time-varying response of a neuron can be predicted by convolution of a linear filter with the sound spectrogram followed by a static rectifying nonlinearity. The LN model is a generalization of the classical spectro-temporal receptive field (STRF), which does not specify a static nonlinearity but provides a similar description of auditory coding (Aertsen and Johannesma 1981; Eggermont et al. 1983; deCharms et al. 1998; Klein et al. 2000; Theunissen et al. 2001). The LN model has been useful because of its generality; that is, because it provides a representation of a neuron’s response properties that holds for arbitrary stimuli (Aertsen and Johannesma 1981; Theunissen et al. 2000).

Despite the relative value of the LN model, it fails to account for important aspects of auditory coding, particularly in more central auditory areas such as primary auditory cortex (A1) (Machens et al. 2004; David et al. 2007; Atencio and Christoph 2008). Several studies have identified nonlinear selectivity for spectro-temporal sound features (Eggermont 1993; Atencio et al. 2008; Sadagopan and Wang 2009; Kozlov and Gentner 2016). In addition, auditory neurons undergo slower changes in response properties that reflect the sensory context (Hiroki and Zador 2009; Rabinowitz et al. 2012). A canonical example of context-dependent changes is stimulus-specific adaptation (SSA), where responses are reduced for repeating versus oddball tones (Ulanovsky et al. 2003). Context-dependent changes in coding are also apparent in responses to clean (undistorted) versus noisy stimuli (Moore et al. 2013; Rabinowitz et al. 2013; Mesgarani et al. 2014). These context effects can last hundreds of milliseconds and reflect nonlinear computations outside of the scope of a linear filter (David and Shamma 2013). Several studies have attempted to bridge this gap in model performance by extending the LN model to incorporate experimentally-observed nonlinearities, including short-term synaptic plasticity (STP) and contrast-dependent gain control (GC) (Rabinowitz et al. 2012; Cooke et al. 2018; Lopez Espejo et al. 2019). These models show improved performance over the LN model and point to mechanisms that explain contextual effects.

The improved performance of these alternatives to the LN model is well established, but the extent to which they describe distinct, complementary mechanisms within the brain is not clear. It has been suggested, for example, that short-term plasticity may in fact contribute to contrast-dependent gain control (Carandini et al. 2002; Rabinowitz et al. 2011). At the same time, both gain control and STP have been implicated in the robust coding of natural stimuli in noise (Rabinowitz et al. 2013; Mesgarani et al. 2014).

The complementarity of these effects has been difficult to establish because thus far they have been tested on datasets recorded from different experimental preparations and using different stimulus sets (Rabinowitz et al. 2012; David and Shamma 2013; Lopez Espejo et al. 2019). To address this issue, we tested both STP and GC models on two natural sounds datasets, collected from A1 of unanesthetized ferret. The first was comprised of a large collection of diverse natural sounds, while the second contained only ferret vocalizations with and without additive broadband noise. We focused on natural sound coding because LN models are limited in their ability to predict responses to natural sounds in A1 (Theunissen et al. 2000; David et al. 2009; Sharpee et al. 2011).

With the models on equal footing, we compared their performance to each other and to a standard LN model. Both models showed improved performance over the LN model, but a model combining the STP and GC mechanisms performed better than either one alone. Additionally, we found a low degree of similarity between the STP and GC models’ predictions after accounting for the LN model’s contributions. These results suggest that models for short-term plasticity and contrast-dependent gain control are not equivalent, and in fact account for complementary components of auditory cortical coding.

## Materials and Methods

### Experimental Procedures

#### Data collection

All procedures were approved by the Oregon Health and Science University Institutional Animal Care and Use Committee (protocol IP00001561) and conform to standards of the Association for Assessment and Accreditation of Laboratory Animal Care (AAALAC).

Prior to experiments, all animals (Mustela putorius furo, 7 males) were implanted with a custom steel head post to allow for stable recording. While under anesthesia (ketamine followed by isoflurane) and under sterile conditions, the skin and muscles on the top of the head were retracted from the central 4 cm diameter of skull. Several stainless steel bone screws (Synthes, 6mm) were attached to the skull, the head post was glued on the mid-line (3M Durelon), and the site was covered with bone cement (Zimmer Palacos). After surgery, the skin around the implant was allowed to heal. Analgesics and antibiotics were administered under veterinary supervision until recovery.

After animals fully recovered from surgery and were habituated to a head-fixed posture, a small craniotomy (1–2 mm diameter) was opened over A1. Neurophysiological activity was recorded using tungsten microelectrodes (1–5 MO, A.M. Systems). One to four electrodes positioned by independent microdrives (Alpha-Omega Engineering EPS) were inserted into the cortex.

Electrophysiological activity was amplified (A.M. Systems 3600), digitized (National Instruments PCI-6259), and recorded using the MANTA open-source data acquisition software (Englitz et al. 2013). Recording site locations were confirmed as being in A1 based on tonotopy, relatively well-defined frequency tuning and short response latency (Kowalski et al. 1996).

Spiking events were extracted from the continuous raw electrophysiological trace by principal components analysis and k-means clustering (David et al. 2009). Single unit isolation was quantified from cluster variance and overlap as the fraction of spikes that were likely to be from a single cell rather than from another cell. Only units with greater than 80% isolation were used for analysis.

Stimulus presentation was controlled by custom software written in Matlab (version 2012A, Mathworks). Digital acoustic signals were transformed to analog (National Instruments PCI6259) and amplified (Crown D-75a) to the desired sound level. Stimuli were presented through a flat-gain, free-field speaker (Manger) 80 cm distant, 0-deg elevation and 30-deg azimuth contralateral to the neurophysiological recording site. Prior to experiments, sound level was calibrated to a standard reference (Brüel & Kjær). Stimuli were presented at 60–65 dB SPL.

#### Natural stimuli

The majority of data included in this study were collected during presentation of a library of natural sounds (set 1: 93, 3 sec/sample, set 2: 306, 4 sec/sample). Some of these sounds (set 1: 30%, set 2: 10%) were ferret vocalizations. The vocalizations were recorded in a sound-attenuating chamber using a commercial digital recorder (44-KHz sampling, Tascam DR-400). Recordings included infant calls (1 week to 1 month of age), adult aggression calls, and adult play calls. No animals that produced the vocalizations in the stimulus library were used in the current study. The remaining natural sounds were drawn from a large library of human speech, music and environmental noises developed to characterize natural sound statistics (McDermott et al. 2013).

Neural activity was recorded during 3 repetitions of these stimuli (set 1: 90, set 2: 288) in random order and either 24 or 30 repetitions of the remaining stimuli (set 1: 3, set 2: 18), all ferret vocalizations, presented on random interleaved trials with 1-3 seconds of silence between stimuli. The low-repetition data were used for model estimation and the high-repetition data were used for model validation.

A second dataset was collected during presentation of ferret vocalizations in clean and noisy conditions. Forty 3-sec vocalizations were each presented without distortion (clean) and with additive Gaussian white noise (0 dB SNR, peak-to-peak). The noise started 0.5 sec prior to the onset and ended 0.5 sec following the offset of each vocalization. A distinct frozen noise sample was paired with each vocalization to allow repetition of identical noisy stimuli. Stimuli were presented at 65 dB SPL with 1-sec inter-stimulus interval.

### Modeling framework

#### Cochlear filterbank

To represent the input for all the encoding models, stimulus waveforms were converted into spectrograms using a second-order gammatone filterbank (Katsiamis et al. 2007). The filterbank included *F* = 18 filters with *f_i_* spaced logarithmically from *f_low_* = 200 to *f_high_* = 20, 000 Hz. After filtering, the signal was smoothed and down-sampled to 100 Hz to match the temporal bin size of the PSTH, and log compression was applied to account for the action of the cochlea.

#### Linear-nonlinear spectrotemporal receptive field (LN) model

The first stage of the LN model applied a finite impulse response (FIR) filter, *h*, to the stimulus spectrogram, *s*, to generate a linear firing rate prediction (*y_lin_*):

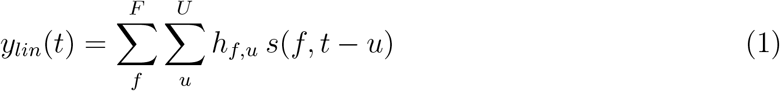

For this study, the filter consisted of *F* = 18 spectral channels and *U* = 15 temporal bins (10ms each). In principle, this step can be applied to the spectrogram as a single 18×15 filter. In practice, the filter was applied in two stages: multiplication by an 18×3 spectral weighting matrix followed by convolution with a 3×15 temporal filter. Previous work has shown that this rank-3 approximation of the full filter is advantageous for prediction accuracy in A1 (Thorson et al. 2015).

The output of the filtering operation was then used as the input to a static sigmoid nonlinearity that mimicked spike threshold and firing rate saturation to produce the final model prediction. For this study, we used a double exponential nonlinearity:

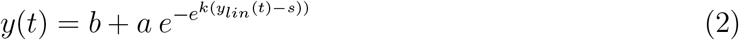

where the baseline spike rate, saturated firing rate, firing threshold, and gain are represented by *b, a, s,* and *k*, respectively.

#### Short-term plasticity (STP) model

The output of each spectral channel served as the input to a virtual synapse that could undergo either depression or facilitation (Tsodyks et al. 1998). In this model, the number of presynaptic vesicles available for release within a virtual synapse is dictated by the fraction of vesicles released by previous stimulation, *u_i_*, and a recovery time constant, *τ_i_*. For depression, *u_i_* > 0, and the fraction of available vesicles, *d*(*t*), is updated,

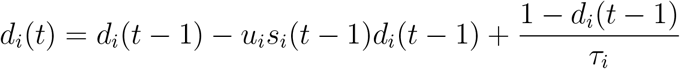

For facilitation, *u_i_* < 0, and *d*(*t*) is updated,

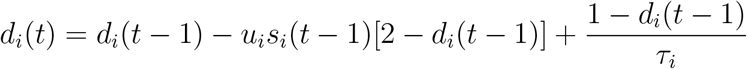

The input to the *i*-th synapse, *s_i_*, is scaled by the fraction of available vesicles, *d_i_*:

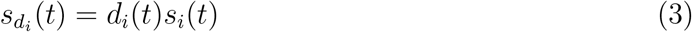

The scaled output, 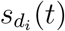, is then used as the input to a temporal filter, identical to the one used in the LN model. Three virtual synapses were used in this study to match the rank-3 STRF approximation, for a total of six free parameters *τ_i_*, *u_i_*, *i* = 0, 1, 2. Values of *τ* and *u* reported in the results represent the mean across all three virtual synapses.

#### Contrast-dependent gain control (GC) model

The contrast-dependent gain control (GC) model was adapted from Rabinowitz et al. 2012. In this model, the parameters of the output nonlinearity depend on a linear weighted sum of the time-varying stimulus contrast. For each stimulus, the contrast, *C*, within a frequency band, *f*, was calculated as the coefficient of variation,

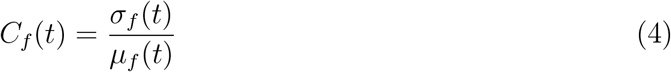

within a 70ms rolling window offset by 20ms. *σ_f_* is the standard deviation and *μ_f_* is the mean level within that window (dB SPL). In the GC model’s original formulation, a linear filter with fittable coefficients would then be applied to the stimulus contrast (Rabinowitz et al. 2012). For this study, we found that a simple contrast weighting provided more accurate predictions. Our implementation used a fixed filter that summed stimulus contrast across frequencies and at a single time point. Thus the contrast at each point reflected the ratio in (4) computed over the window 20-90 ms preceding the current moment in time. The output, *K*, of this summation was then used to determine the parameters of the output nonlinearity in (2) such that the *i*-th parameter, *θ_i_*, was determined from the base value, 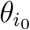, that would normally be fitted in (2) and a contrast weighting term, 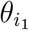 :

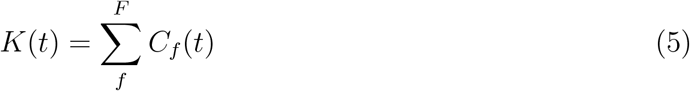

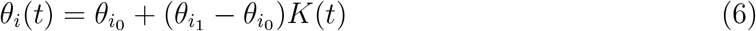

With this formula, it’s necessary to know both 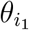 and 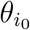 to determine the impact of contrast on any particular parameter. However, we did not find any significant differences in the base values 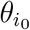 between improved and non-improved cells. In the results we instead report the difference, 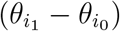, which represents the slope of the linear relationship proposed by the model.

#### Model optimization

Models were optimized using the L-BFGS-B gradient descent algorithm implemented in the SciPy Python library (Virtanen et al. 2020). This optimization minimized the mean-squared error (MSE) between a neuron’s time-varying response, averaged across any repeated presentations of the same stimulus, and the model’s prediction. Post-fitting performance was evaluated based on the correlation coefficient (Pearson’s R) between prediction and response, adjusted for the finite sampling of validation data (Hsu et al. 2004).

#### Equivalence analysis

Equivalence of STP and GC models was quantified using the partial correlation between the time-varying response to the validation stimuli predicted by each model, computed relative to the prediction by the LN model (Baba et al. 2004). If the two models were equivalent and deviated from the LN model in exactly the same way, the partial correlation would be 1.0. If they deviated in completely different ways, the partial correlation would be 0.0.

Because data set size was finite, noise in the estimation data produced uncertainty in model parameter estimates, which biased partial correlation toward values less than 1. To compute an upper bound on equivalence, we split the estimation data set in half and fit the same model (STP or GC) to both halves. We then measured equivalence between the same model fit to each half. This upper bound was itself corrected to account for the fact that each model was fit with only half the estimation data. We applied a scaling factor to the median equivalence score to account for the effect of using a smaller dataset for fitting.

The scaling factor was the ratio of the median partial correlation between-models for the full estimation data (0.204) to the median partial correlation between-models for opposite halves of the estimation data (0.123) on the subset of cells for which the within-model analysis was performed, resulting in a scaling factor of 1.66.

#### Software Accessibility

The software described in the paper is freely available online at https://github.com/LBHB/NEMS and https://github.com/LBHB/nems_db. Analyses for this study were run on a private compute cluster using Intel core-i7 CPUs and the Ubuntu operating system version 16.04.

## Results

### Models for encoding of natural sounds by neurons in primary auditory cortex

To compare performance of the short-term plasticity (STP) and contrast-dependent gain control (GC) models directly, we recorded the activity of *n* = 540 neurons in A1 of awake, non-behaving ferrets during presentation of a large natural sound library (Fig. 1). We then compared how models incorporating STP or GC nonlinearities accounted for sound-evoked activity in the same dataset. See Table 1 for a list of all statistical tests used, with individual tests referenced by superscript throughout the results.

**Table 1:** Statistical tests reported in the Results, labeled in the text by the letters in the left-hand column. *p*^∗^ indicates Bonferroni-adjusted *p* values for 12 multiple comparisons.

|  | Distribution | Test | Statistic | P-value | Sample Size | Animals |
| --- | --- | --- | --- | --- | --- | --- |
| a | Non-parametric | Permutation Test | N/A | $p < 0.05$, 468 sig. | $n = 540$ | $n = 7$ |
| b | Non-parametric | Wilcoxon Signed Rank | $T = 2.17 \times 10^4$ | $p = 7.54 \times 10^{-30}$ | $n = 468$ | $n = 7$ |
| c | Non-parametric | Wilcoxon Signed Rank | $T = 2.39 \times 10^4$ | $p = 3.55 \times 10^{-26}$ | $n = 468$ | $n = 7$ |
| d | Non-parametric | Wilcoxon Signed Rank | $T = 4.46 \times 10^4$ | $p = 5.00 \times 10^{-4}$ | $n = 468$ | $n = 7$ |
| e | Non-parametric | Wilcoxon Signed Rank | $T = 2.60 \times 10^4$ | $p = 5.10 \times 10^{-23}$ | $n = 468$ | $n = 7$ |
| f | Non-parametric | Wilcoxon Signed Rank | $T = 2.31 \times 10^4$ | $p = 2.02 \times 10^{-27}$ | $n = 468$ | $n = 7$ |
| g | Non-parametric | Jackknife T-test | T (varies) | $p < 0.05$, 132 sig. | $n = 468$ | $n = 7$ |
| h | Normal (residuals) | Pearson's Correlation | $r = 0.18$ | $p = 6.42 \times 10^{-5}$ | $n = 468$ | $n = 7$ |
| i | Normal (residuals) | Pearson's Correlation | $r = 0.31$ | $p = 1.26 \times 10^{-6}$ | $n = 237$ | $n = 2$ |
| j | Normal (residuals) | Pearson's Correlation | $r = 0.50$ | $p = 1.34 \times 10^{-16}$ | $n = 237$ | $n = 2$ |
| k | Non-parametric | Mann-Whitney U | $U = 1.48 \times 10^4$ | $p = 1.72 \times 10^{-8}$ | $n = 132, 336$ | $n = 7$ |
| l | Normal (residuals) | Pearson's Correlation | $r = 0.22$ | $p = 0.0103$ | $n = 132$ | $n = 7$ |
| m | Non-parametric | Wilcoxon Signed Rank | $T = 4.90 \times 10^4$ | $p = 0.0444$ | $n = 468$ | $n = 7$ |
| n | Non-parametric | Mann-Whitney U | $U = 1.61 \times 10^4$ | $p = 3.53 \times 10^{-6}$ | $n = 132, 336$ | $n = 7$ |
| o | Non-parametric | Mann-Whitney U | $U = 1.26 \times 10^4$ | $p = 2.92 \times 10^{-13}$ | $n = 132, 336$ | $n = 7$ |
| p | Non-parametric | Mann-Whitney U | $U = 2.72 \times 10^4$ | $p = 1.22 \times 10^{-4}$ | $n = 132, 336$ | $n = 7$ |
| q | Non-parametric | Mann-Whitney U | $U = 2.81 \times 10^4$ | $p = 8.01 \times 10^{-6}$ | $n = 132, 336$ | $n = 7$ |
| r | Non-parametric | Mann-Whitney U | $U = 1.56 \times 10^4$ | $p = 5.90 \times 10^{-7}$ | $n = 132, 336$ | $n = 7$ |
| s | Non-parametric | Mann-Whitney U | $U = 2.24 \times 10^4$ | $p = 0.852 \times 10^{-4}$ | $n = 132, 336$ | $n = 7$ |
| t | Non-Parametric | Mann-Whitney U | $U = 1.94 \times 10^3$ | $p^* = 1.00$ | $n = 132, 336$ | $n = 7$ |
| u | Non-Parametric | Mann-Whitney U | $U = 5.19 \times 10^2$ | $p^* = 1.00$ | $n = 132, 336$ | $n = 7$ |
| v | Non-Parametric | Mann-Whitney U | $U = 5.28 \times 10^2$ | $p^* = 1.00$ | $n = 132, 336$ | $n = 7$ |
| w | Non-Parametric | Mann-Whitney U | $U = 3.39 \times 10^3$ | $p^* = 1.00$ | $n = 132, 336$ | $n = 7$ |
| x | Non-Parametric | Mann-Whitney U | $U = 1.09 \times 10^4$ | $p^* = 1.00$ | $n = 132, 336$ | $n = 7$ |
| y | Non-Parametric | Mann-Whitney U | $U = 9.49 \times 10^3$ | $p^* = 1.00$ | $n = 132, 336$ | $n = 7$ |
| z | Non-Parametric | Mann-Whitney U | $U = 1.86 \times 10^3$ | $p^* = 1.00$ | $n = 132, 336$ | $n = 7$ |
| aa | Non-Parametric | Mann-Whitney U | $U = 5.47 \times 10^2$ | $p^* = 1.00$ | $n = 132, 336$ | $n = 7$ |
| bb | Non-Parametric | Mann-Whitney U | $U = 2.37 \times 10^4$ | $p^* = 1.00$ | $n = 132, 336$ | $n = 7$ |
| cc | Non-Parametric | Mann-Whitney U | $U = 4.78 \times 10^2$ | $p^* = 1.00$ | $n = 132, 336$ | $n = 7$ |
| dd | Non-Parametric | Mann-Whitney U | $U = 1.40 \times 10^4$ | $p^* = 1.68 \times 10^{-3}$ | $n = 132, 336$ | $n = 7$ |
| ee | Non-Parametric | Mann-Whitney U | $U = 1.19 \times 10^4$ | $p^* = 2.87 \times 10^{-3}$ | $n = 132, 336$ | $n = 7$ |
| ff | Non-parametric | Wilcoxon Signed Rank | $T = 3.67 \times 10^3$ | $p = 5.84 \times 10^{-3}$ | $n = 141$ | $n = 6$ |
| gg | Non-parametric | Wilcoxon Signed Rank | $T = 2.26 \times 10^3$ | $p = 1.70 \times 10^{-8}$ | $n = 141$ | $n = 6$ |
| hh | Non-parametric | Wilcoxon Signed Rank | $T = 2.14 \times 10^3$ | $p = 3.69 \times 10^{-9}$ | $n = 141$ | $n = 6$ |
| ii | Non-parametric | Wilcoxon Signed Rank | $T = 3.02 \times 10^3$ | $p = 4.27 \times 10^{-5}$ | $n = 141$ | $n = 6$ |
| jj | Non-parametric | Wilcoxon Signed Rank | $T = 4.22 \times 10^3$ | $p = 0.105$ | $n = 141$ | $n = 6$ |
| kk | Normal (residuals) | Pearson's Correlation | $r = 0.18$ | $p = 0.0345$ | $n = 141$ | $n = 6$ |
| ll | Non-parametric | Mann-Whitney U | $U = 5.02 \times 10^2$ | $p = 0.0449$ | $n = 12, 129$ | $n = 6$ |

**Figure 1:**
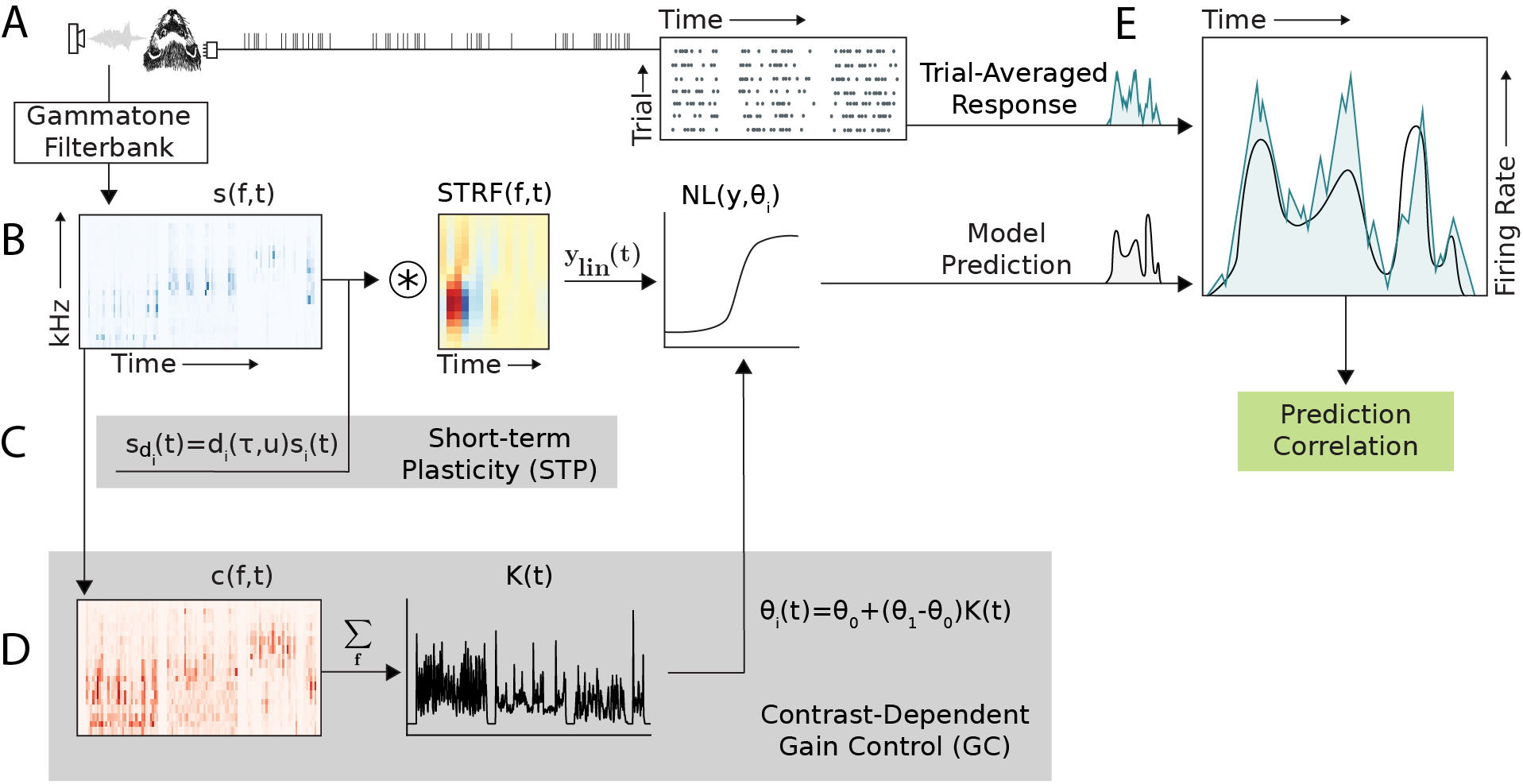
Schematic of four different architectures for modeling sound encoding by neurons in auditory cortex. (A) Single neuron activity was recorded from primary auditory cortex (A1) of awake, passively listening ferrets during presentation of a large set of natural sound stimuli. The trial-averaged response to each sound was calculated as the instantaneous firing rate using 10ms bins. Sound waveforms were transformed into 18-channel spectrograms with log-spaced frequencies for input to the models. (B) Linear nonlinear spectro-temporal receptive field (LN) model: stimulus spectrogram is convolved with a linear spectro-temporal filter followed by nonlinear rectification. (C) Short-term plasticity (STP) model: simulated synapses depress or facilitate spectral stimulus channels prior to temporal convolution. (D) Contrast-dependent gain control (GC) model: the coefficient of variation (contrast) of the stimulus spectrogram within a rolling window is summed across frequencies. Parameters for the nonlinear rectifier are scaled by time-varying contrast. (E) Model performance is measured by the correlation coefficient (Pearson’s *R*) between the trial-averaged response and the model prediction. The four architectures were defined as follows - LN: B only, STP: B and C, GC: B and D, GC+STP: B, C, and D.

We compared performance of four model architectures, fitting and evaluating each with the same data set (Fig. 1b-d). The first was a standard LN model, which is widely used to characterize spectro-temporal sound encoding properties (Simoncelli et al. 2003; Calabrese et al. 2011; Rahman et al. 2019) and provided a baseline for the current study. The second architecture (STP model) accounted for synaptic depression or facilitation by scaling input stimuli through simulated plastic synapses (Tsodyks et al. 1998; Wehr and Zador 2005). The third (GC model) scaled a neuron’s sound-evoked spike rate as a function of recent stimulus contrast (Rabinowitz et al. 2011). A fourth architecture (GC+STP model) incorporated both STP and GC mechanisms into the LN model. The LN, STP and GC models were implemented following previously published architectures (Rabinowitz et al. 2012; Lopez Espejo et al. 2019), and the GC+STP model combined elements from the other models in a single architecture. Performance was evaluated using a reserved validation set that was not used for fitting.

### Complementary explanatory power by short-term plasticity and gain control models

We quantified prediction accuracy using the correlation coefficient (Pearson’s *R*) between the predicted and actual peri-stimulus time histogram (PSTH) response in each neuron’s validation data. Before comparing performance between models, we identified auditory-responsive neurons for which the prediction accuracy of all four models was greater than expected for a random prediction (*p* < 0.05, permutation test, *n* = 468/540 ^*a*^). Comparisons then focused on this subset. This conservative choice ensured that comparisons between model performance were not disproportionately impacted by cells for which a particular optimization failed.

A comparison of median prediction correlation across the entire set of auditory-responsive neurons (*n* = 468, Fig. 2a) revealed that both the GC model and the STP model performed significantly better than the LN model (*p* = 7.54 × 10^−30^ and *p* = 3.55 × 10^−26^, respectively ^*b,c*^, two-sided Wilcoxon signed-rank test), confirming previous results (Rabinowitz et al. 2012; Lopez Espejo et al. 2019). We also found that the STP model performed significantly better than the GC model (*p* = 5.00 × 10^−24 *d*^). If the STP and GC models were equivalent to one another, we would not expect to observe further improvement for the combined GC+STP model. Instead, we observed a significant increase in predictive power for the combined model over both the GC model and the STP model (*p* = 5.10 × 10^−^23 and *p* = 2.02 × 10^−^27, respectively ^*e,f*^). The improvement for the combined model suggests that the STP and GC models describe complementary functional properties of A1 neurons.

**Figure 2:**
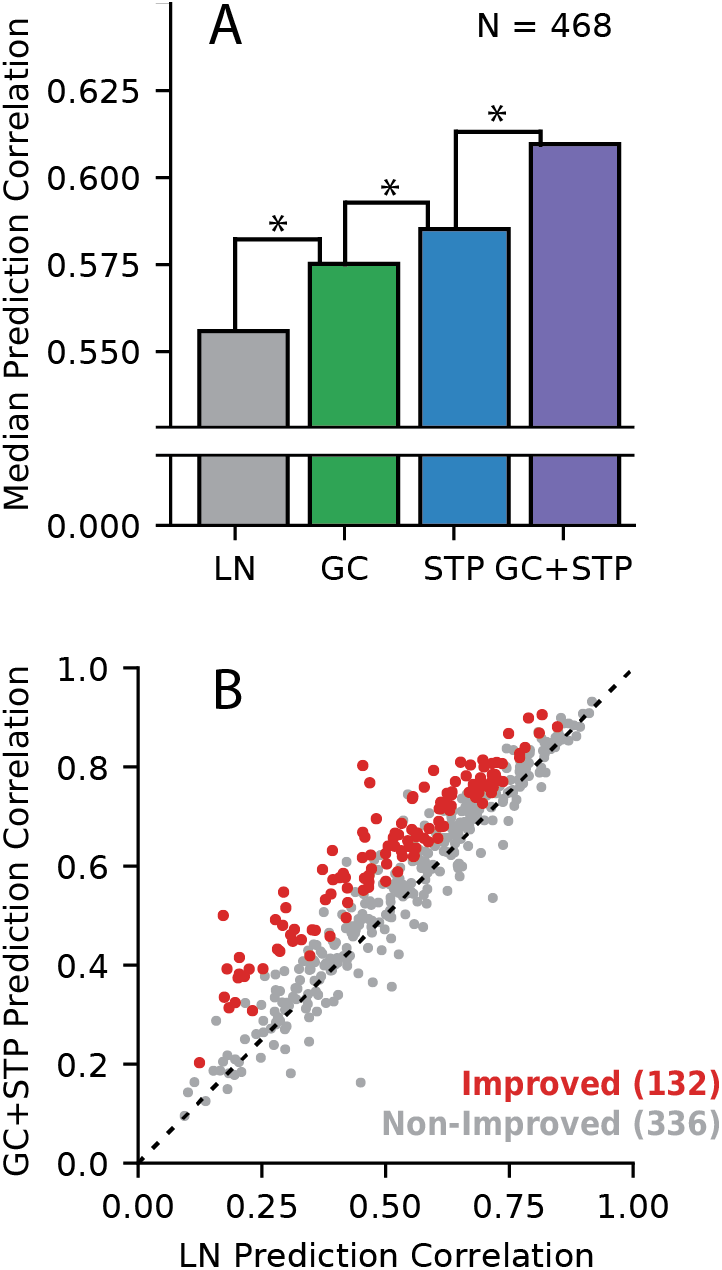
Comparison of model prediction accuracy. (A) Median prediction correlation for each model (*n* = 468 neurons). Differences between LN & GC (*p* = 7.54 × 10^−30^), GC & STP (*p* = 5.00 × 10^−4^), and STP & GC+STP (*p* = 2.02 × 10^−27^) models were all significant (, ∗ < 0.05, two-sided Wilcoxon signed-rank test). (B) Scatter plot compares prediction correlation by the LN model and combined GC+STP model for each neuron. Color indicates whether the combined model showed a significant improvement (red, *p* < 0.05, permutation test) or not (gray).

The scatter plot in Figure 2b compares performance of the LN and combined models for each neuron. Among the 468 auditory-responsive neurons, 132 (28.2%) showed a significant improvement in prediction accuracy for the combined versus the LN model (*p* < 0.05, jackknife t-test, ^*g*^). For the analyses of model equivalence and parameter distributions below, we focus on this set of improved neurons.

### Limited equivalence of short-term plasticity and contrast gain model predictions

A central question in this study was the extent to which the STP model’s improved performance over the LN model could be accounted for by the GC model, or vice-versa. Among non-improved cells, the two models’ predictions were often closely matched to each other and to the prediction of the LN model (Fig. 3a). However, for some improved neurons, the time-varying responses predicted by the STP and GC models were readily distinguishable not only from the LN model but also from each other (Fig. 3b,c). In this case, the STP and GC models both improved prediction accuracy, but they did so with low equivalence. That is, the models’ predicted responses deviated from that of the LN model in different ways. If the STP and GC models accounted for equivalent nonlinear properties, their predicted responses should remain similar to each other, even when differing from the LN model.

**Figure 3:**
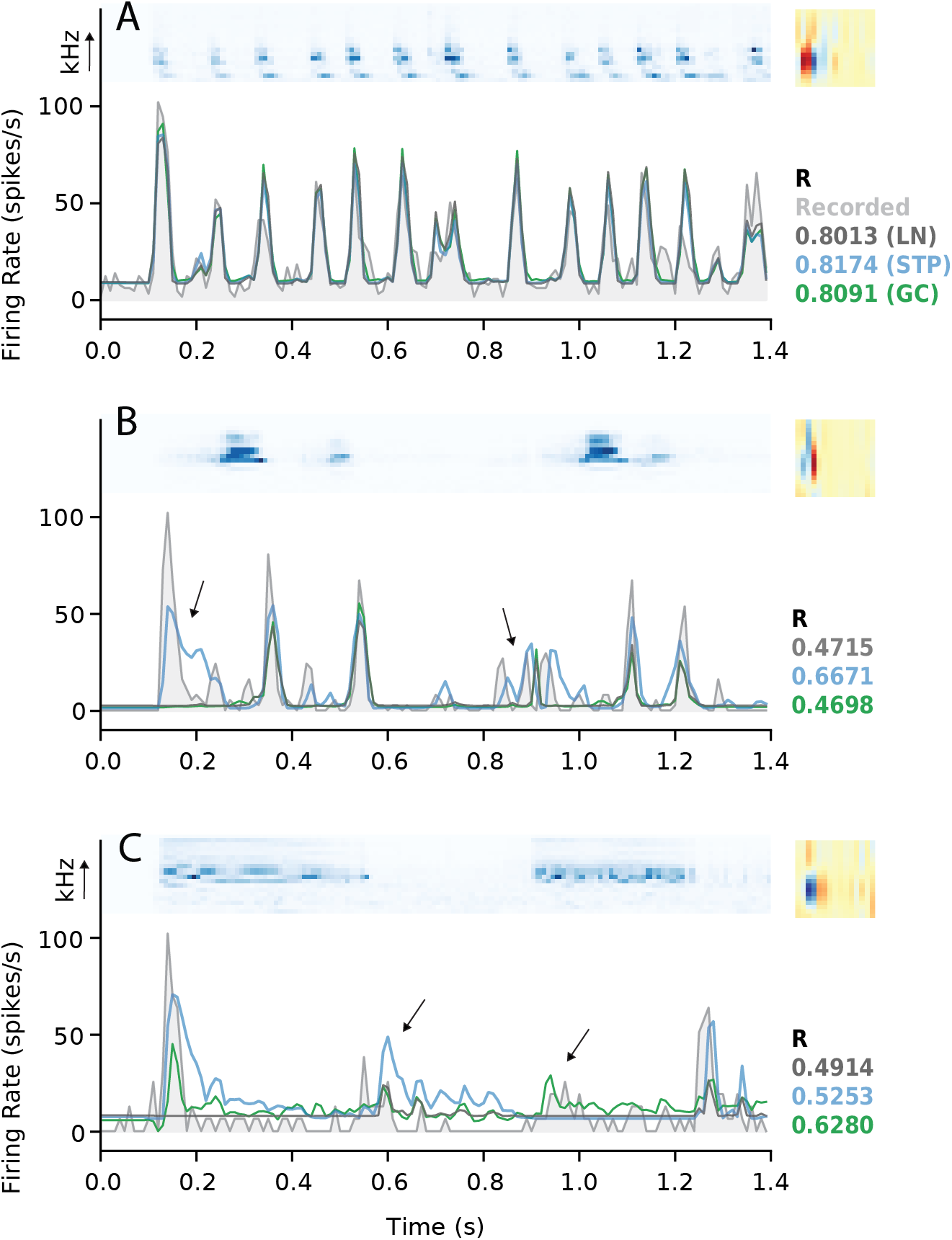
Example model fits and predictions. (A) Results from a neuron for which the STP and GC model predictions were not significantly better than the LN model prediction. Top left subpanel shows the spectrogram from one natural sound in the validation set. Top right panel shows the spectro-temporal filter from the LN model fit (right). Bottom panel shows the actual PSTH response (light gray, filled) overlaid with predictions by the LN (dark gray), STP (blue), and GC (green) models. Values at the bottom right of each panel indicate the prediction correlations for each model in the corresponding color. The actual response was smoothed using a 30ms boxcar filter for visualization. (B) Data from a neuron for which the STP model performed significantly better than the LN and GC models, plotted as in A. Arrows indicate times for which the STP model successfully reproduced an increase in firing rate while the other models did not. (C) Data from a neuron for which the GC model performed significantly better than the LN and STP models. Arrows indicate a time for which the STP model incorrectly predicted an increase in firing rate (left) and a time for which the GC model successfully reproduced an increase in firing rate while the other models did not (right).

To quantify model equivalence across all neurons, we first compared the change in prediction correlation for the STP and GC models, relative to the LN model (Fig. 4a). If the two models were equivalent, we would expect a strong positive correlation between improvements over the LN model per neuron. However, we observed only a weak correlation (r = 0.18, *p* = 6.42 × 10^−5 *h*^). To estimate the expected strength of such a relationship for models known to be equivalent, we performed a within-model comparison on a subset of cells for which responses to a larger collection of sounds were recorded (*n* = 237). We fit the LN, STP, and GC models twice, each time using a separate half of the estimation data. The within-model correlation between changes in prediction correlation, relative to the LN model fit on the same half of the estimation set, was significantly higher for both the STP and GC models (*r* = 0.50, *p* = 1.34 × 10^−16^ and *r* = 0.31, *p* = 1.26 × 10^−6^, respectively^*i,j*^).

**Figure 4:**
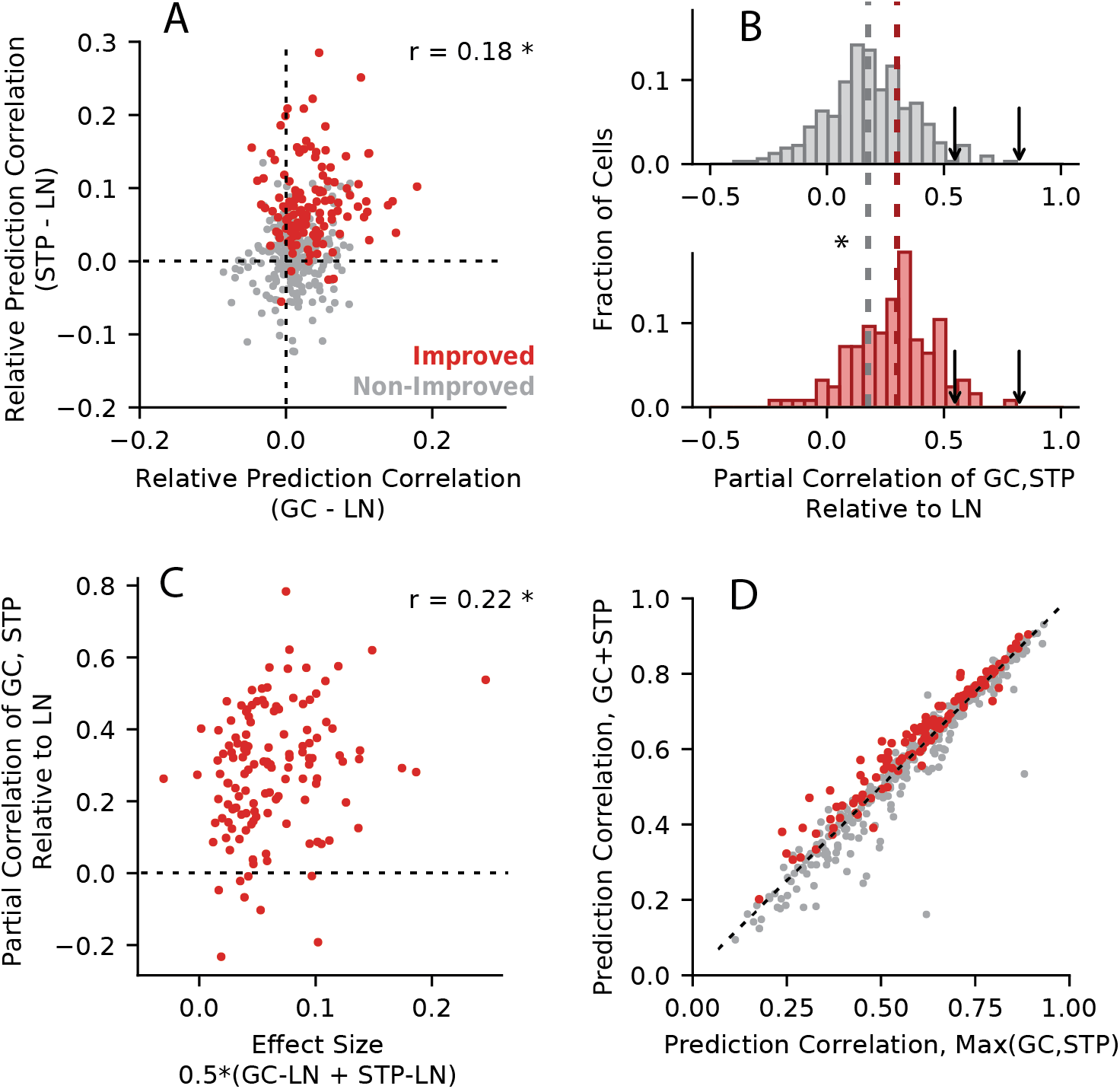
Equivalence of model predictions. (A) Difference in prediction correlation between the GC (horizontal axis) or STP (vertical axis) model and the LN model for each neuron (*r* = 0.18, *p* = 6.42 × 10^−5^). Red points indicate neurons with a significant improvement for the GC+STP model over the LN model (*p* < 0.05, permutation test); gray points indicate neurons that were not improved. (B) Histogram of model equivalence for each unit, measured as the partial correlation between time-varying response predicted by the STP and GC models relative to the LN model prediction. Median equivalence for improved cells (0.29, bottom, red) was significantly greater than for non-improved cells (0.17, top, gray) (Mann-Whitney U test, *p* = 1.72 × 10^−8^). Arrows indicate median partial correlation for the GC model (0.54, left) and the STP model (0.81, right) when compared within-model, adjusted for differences in estimation data. (C) Scatter plot compares equivalence (vertical axis) versus effect size (horizontal axis), *i.e.*, the average change in prediction correlation for the STP and GC models relative to the LN model. Only a weak relationship between equivalence and effect size was observed (*r* = 0.22, *p* = 0.0103). (D) Prediction correlations for the combined GC+STP model (vertical axis) and the maximum of the GC and STP models (horizontal axis). The median prediction correlation for the combined model (0.6075) was significantly smaller than the greater of the individual models (0.6088), though the difference in medians was quite small (Wilcoxon signed rank test, *p* = 0.0444)

If two models account for the same functional properties, we also expect them to predict the same time-varying response to a test stimulus. To test for this possibility, we measured the similarity of responses predicted by the STP and GC models. Since both models are extensions of the LN model, we discounted the contributions of the LN model to the prediction (Fig. 4b). We defined model equivalence for each neuron as the partial correlation between the STP and GC model predictions relative to the LN model prediction (Baba et al. 2004). An equivalence score of 1.0 would indicate perfectly equivalent STP and GC model predictions, and a score of 0 would indicate that the models accounted for completely different response properties.

For *n* = 132 neurons with improvements over the LN model, the distribution of partial correlations had a median of 0.29. This value was relatively low, again indicating only weak equivalence between the models. However, it was significantly greater than the median for non-improved neurons (0.17) (*p* = 1.72 × 10^−8^, Mann-Whitney U test ^*k*^). Thus, there are some similarities between the neural dynamics accounted for by the STP and GC models. Because of estimation error, one would expect measured equivalence to be less than the theoretical maximum of 1.0. To quantify the upper bound on partial correlation, we measured equivalence between two fits of the same model, using the same split estimation data described above for the relative performance comparison. We scaled these values according to the amount of noise introduced by the reduced size of the split estimation dataset to get adjusted medians of 0.81 for the STP model and 0.54 for the GC model. Thus, while the STP and GC models show some degree of equivalence, it is lower than would be expected for fully equivalent models.

We performed one additional control for the possibility that low equivalence scores resulted from model estimation noise. If noise was the major factor, then equivalence should be higher for models with better prediction accuracy. For some cells, improvements for the STP and GC models were small (*e.g.*, Fig. 3a). If the deviations from the LN model prediction were small and mostly reflected noise, one would expect weak equivalence. We therefore defined a measure of effect size for each cell as the mean change in prediction correlation for the STP and GC models relative to the LN model. Neurons for which the more complex models did not improve prediction accuracy compared to the LN model should have a low equivalence score, but would also have a small effect size. If the STP and GC models equivalent, we would expect most cells with significant improvements over the LN model to only have a large effect size if they also had a high equivalence score. However, we did not discern any clear pattern in the data (Fig. 4c). Instead, equivalence and effect size were only weakly related (*r* = 0.22, *p* = 0.0103 ^*l*^).

Following this evidence for limited equivalence, we asked whether the combined model’s greater predictive power was merely the result of its ability to simply account for either STP or GC, without any benefit from their combination in individual neurons. If this were the case, we would expect that for any given cell, the prediction correlation of the combined model should be no greater than the larger of the prediction correlations for the STP and GC models (Fig. 4d). Indeed, we found that the median prediction correlation was significantly smaller for the combined model, although the difference was small (median difference 0.0013, Wilcoxon signed rank test, two-sided, *p* = 0.0444 ^*m*^). However, we also observed that prediction accuracy was greater for the combined model than for either the STP or GC model individually for 108 of the 132 improved cells (0.82%). Thus, a small number of neurons exhibit dynamics explained better by a combination of the STP and GC nonlinearities. However, a majority of the non-improved neurons appeared to utilize only one mechanism or the other (189/336, or 56%).

### Simulations highlight distinct functional properties of short-term plasticity and contrast-dependent gain control models

The weak equivalence of the GC and STP models (Fig. 4) suggests that these models account for functionally distinct response properties in A1. As one final control for the possibility that estimation noise could have biased our equivalence results, we analyzed data simulated using the different models. We fit the LN, STP and GC models to three simulated neural responses (Fig. 5). Each simulation was generated by using a fitted model to predict responses to a set of natural stimuli and treating the model’s prediction as ground-truth for subsequent fitting. Because the simulated responses could be generated noise-free, any difference in STP vs. GC model performance could be attributed to differences in their ability account for nonlinear response properties. The first simulated response was generated using the LN model fit to a cell that showed no improvements for the STP or GC models (Fig. 3a). The other simulations were generated using the STP and GC model fits from neurons for which the STP or CG models, respectively, performed better than the LN model (shown in Fig. 3b and 3c).

**Figure 5:**
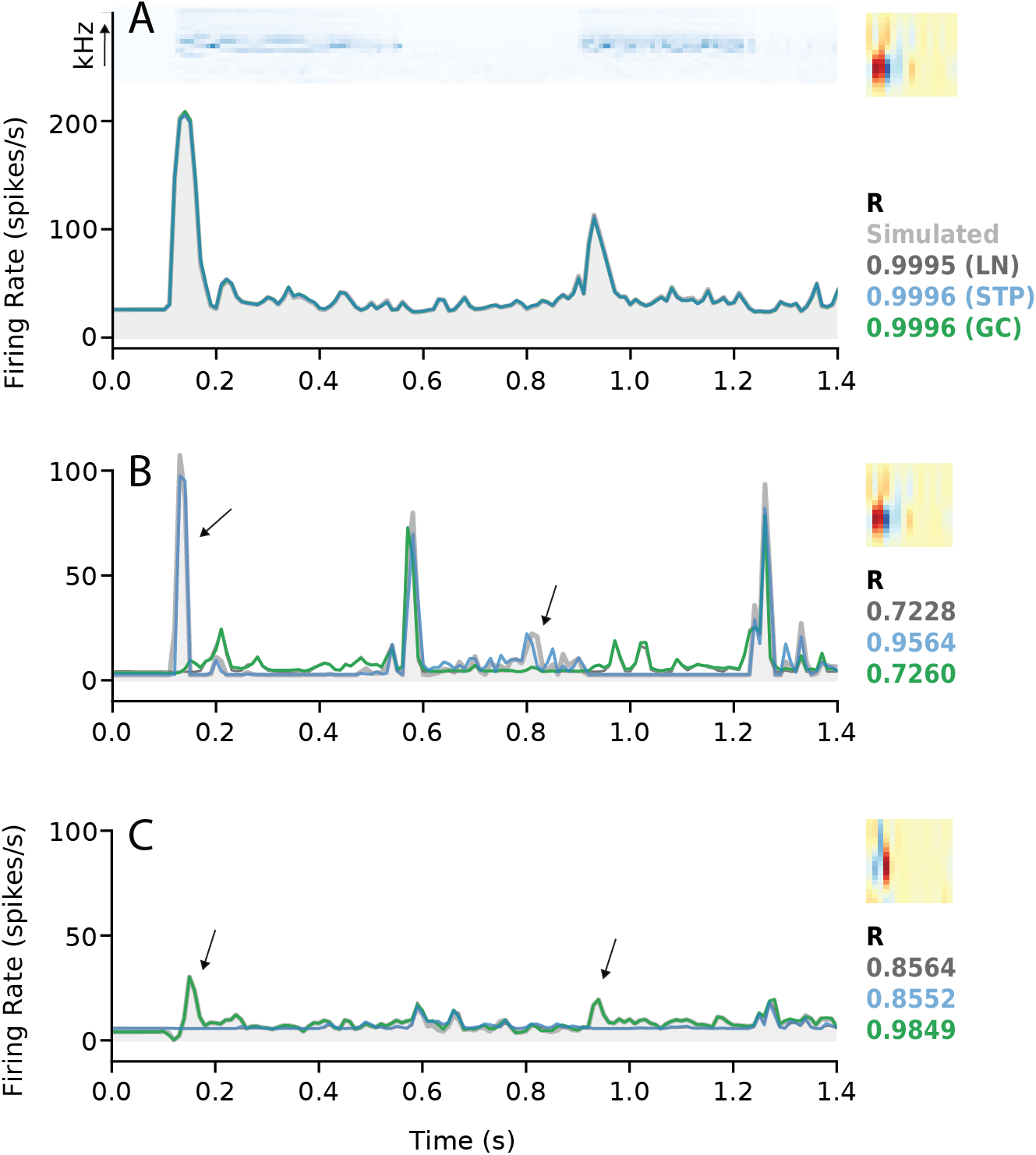
Model performance for simulated data. (A) Simulation based on the fitted LN model from Fig. 3a. Simulated PSTH response to one stimulus (spectrogram at top) based on an LN model is plotted in gray shading. Predicted PSTHs for each model (LN, STP or GC) are overlaid, and model prediction correlation is indicated at right (LN: dark gray, STP: blue, GC: green). For this linear neuron, all three models perform nearly identically. The linear filter fit using the LN model is shown at the top right. (B) Model fits for simulation based on the STP model from Fig. 3b, plotted as in a. (C) Model fits for simulation based on the GC model from Fig. 3c.

As expected, all three models were able to reproduce the LN simulation nearly perfectly (Pearson’s R = 0.9995, 0.9996, and 0.9996 for the LN, STP, and GC models, respectively, Fig. 5a). However, when fit to the STP simulation, the GC model was no better than the LN model, but the STP model did perform better (R = 0.7228, 0.9564, and 0.7260, Fig. 5b). Conversely, when fit to the GC simulation, the STP model performed no better than the LN model while the GC model did (R = 0.8564, 0.8552, 0.9849, Fig. 5c). This pattern of different performance confirmed that that the STP and GC models did account for distinct neuronal dynamics.

### Model fit parameters are consistent with previous studies

Since both the STP model and the GC model used in this study were designed to replicate previous studies (Rabinowitz et al. 2012; Lopez Espejo et al. 2019), it’s important to verify that the models behaved consistently with these previous observations. This consideration was of particular relevance for the GC model since it had not previously been fit using a natural sound dataset. To test for consistency, we analyzed the distributions of their fitted parameter values for comparison with the previous reports.

For the short-term plasticity model (Fig. 6), we found that the median values of both the time constant (*τ*) and fraction gain change (*u*) parameters were significantly higher for improved versus non-improved cells (Mann-Whitney U tests, two-sided, *p* = 3.53 × 10^−6^ and *p* = 2.92 × 10^−13^, respectively ^*n,o*^). The difference was more pronounced for the *u* parameter, and nearly all cells had positive values for the *u* parameter. The large impact of the *u* parameter and predominance of depression over facilitation agreed with published results (Lopez Espejo et al. 2019).

**Figure 6:**
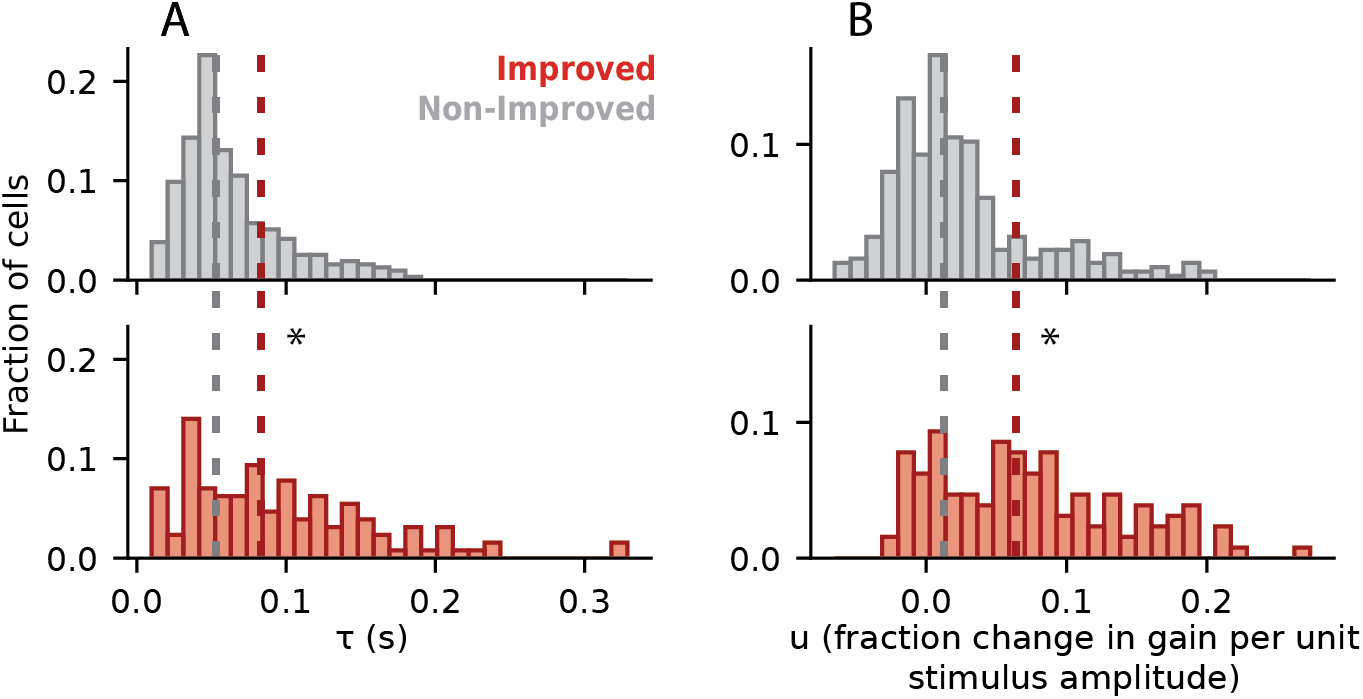
Parameter fit values for the short-term plasticity model. (A) Distribution of *τ*, representing the time constant for the recovery of synaptic vesicles Top panel shows data for non-improved neurons (gray) and bottom panel for improved neurons (red). Median values for non-improved (0.0533 s) and improved (0.0833 s) neurons were significantly different (*p* = 3.53 × 10^−6^, two-sided Mann-Whitney U test, * *p* < 0.05), indicating a longer time constant for the improved cells. (B) Distribution of *u* values, representing release probability (*i.e.*, the fraction change in gain per unit of stimulus amplitude). Medians for non-improved (0.0128) and improved (0.0641) neurons were significantly different (*p* = 2.92 × 10^−13^), showing higher release probability for neurons with improved performance over the LN model.

For the contrast-dependent gain control model (Fig. 7), the impact of contrast on the slope (*k*) parameter of the output nonlinearity was significantly more negative for improved neurons than for non-improved neurons (Mann-Whitney U tests, two-sided; *p* = 1.22 × 10^−4 *p*^). This indicates a decrease in neural gain during high contrast sounds, which is consistent with models fit using dynamic random chord (RC-DRC) stimuli (Rabinowitz et al. 2012). Additionally, compared to non-improved cells, the relationship between contrast and saturation (*a*) was significantly more negative and the relationship between contrast and baseline (*b*) was significantly more positive for improved cells (*p* = 8.01 × 10^−6^ and *p* = 5.90 × 10^−7^, respectively ^*q,r*^). However, we observed no significant difference in the effect of contrast for the input offset (*s*) parameter, which did change for RC-DRC stimuli (*p* = 0.852 ^*s*^). As observed in the previous study, the net result of increasing contrast was to decrease the gain of neural responses, and thus the overall effects of changing contrast on response gain were consistent between studies.

**Figure 7:**
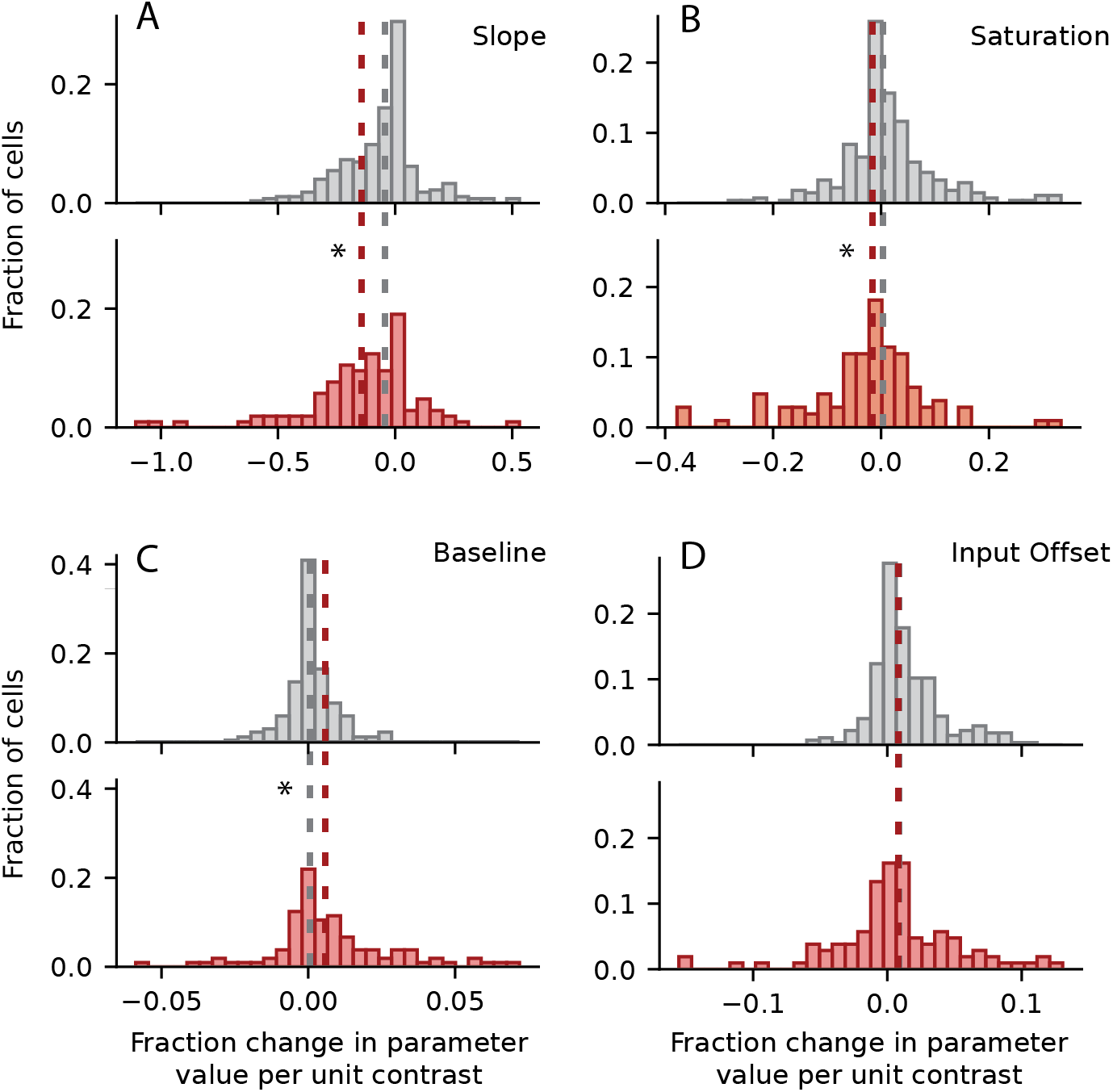
Parameter fit values for the contrast-dependent gain control model. (A) Histogram of effect of contrast on *k*, representing the slope of the output nonlinearity, plotted for non-improved neurons (gray, top) and improved neurons (red, bottom) as in Fig. 6. The negative median value for improved (−0.14) cells indicates a decrease in slope during high-contrast conditions. This median was significantly smaller than for non-improved neurons (−0.042, *p* = 1.22 × 10^−4^, two-sided Mann-Whitney U test). Asterisk (*) indicates *p* < 0.05. (B) Histogram of contrast effect on *a* (saturation level), plotted as in A. Median values for non-improved (0.0031) and improved (−0.0156) neurons were significantly different, indicating a decreased response amplitude in high contrast conditions (*p* = 8.01 × 10^−6^). (C) Histograms of contrast effect on *b* (baseline of the output nonlinearity). Medians for non-improved (0.0005) and improved (0.0058) neurons were significantly different, indicating an increase in baseline for high contrast (*p* = 5.90 × 10^−7^). (D) Distribution of contrast effect on *s* (input offset). There was no significant difference between medians for non-improved (0.0088) and improved (0.0082) neurons (*p* = 0.85).

### Relationship between baseline neural properties and model performance

Previous work has reported that spontaneous firing rates are related to the predictive power of the STP model (David and Shamma 2013). We therefore asked whether there was a basic functional property that could predict whether a particular neuron would benefit from one nonlinear model or another. While we could not distinguish neuronal cell type (e.g., excitatory versus inhibitory neurons), we could measure basic aspects of spiking activity, namely spontaneous and evoked firing rates, that might correspond to biological properties of the neurons. To this end, we split the neurons into four mutually exclusive groups. The first three consisted of neurons for which none of the nonlinear models significantly improved prediction accuracy (“None”, *n* = 327), neurons for which the STP model significantly improved prediction accuracy (“GC”, *n* = 66), and neurons for which the GC model significantly improved prediction accuracy (“STP”, *n* = 17), respectively. The fourth group contained neurons for which both the STP and GC models significantly improved prediction accuracy or for which the *GC* + *STP* model alone significantly improved prediction accuracy (“Both”, *n* = 56). We then separately compared the median evoked and spontaneous firing rates between each group 8.

Of the twelve comparisons made, most (10/12) were not statistically significant significant (*p*^∗^ > 0.05 ^*t*−*cc*^). Only two showed a significant difference: median evoked firing rate was significantly higher for the “STP” (11.7 spikes/s) and “Both” (12.7 spikes/s) groups compared to the “None” (7.31 spikes/s) group (Mann-Whitney U test, *p*^∗^ = 1.68 × 10^−3^ and *p*^∗^ = 2.87 × 10^−3^, respectively ^*dd,ee*^, adjusted for multiple comparisons). Thus, there was a small correlation between evoked activity and STP effects.

### Greater relative contribution of contrast gain to encoding of noisy natural sounds

We also compared performance of the STP and GC models on a smaller data set collected with clean and noisy ferret vocalizations 9. Previous studies using stimulus reconstruction methods argued that both short-term plasticity and contrast-dependent gain control are necessary for robust encoding of noisy natural signals (Rabinowitz et al. 2013; Mesgarani et al. 2014). The inclusion of additive noise should reduce contrast by increasing the mean sound energy and reducing variance. We compared the mean and standard deviation of each sample for the two data sets to see if there were any systematic differences in contrast. We found that the natural sounds dataset smoothly spanned the range of observed means and standard deviations. Meanwhile, the vocalization dataset formed two distinct categories categories: a high-contrast group of clean vocalizations and a low-contrast group of noisy vocalizations (Fig. 10). A small set of the natural sounds overlapped with the noisy vocalizations because they had similar noise characteristics.

**Figure 8:**
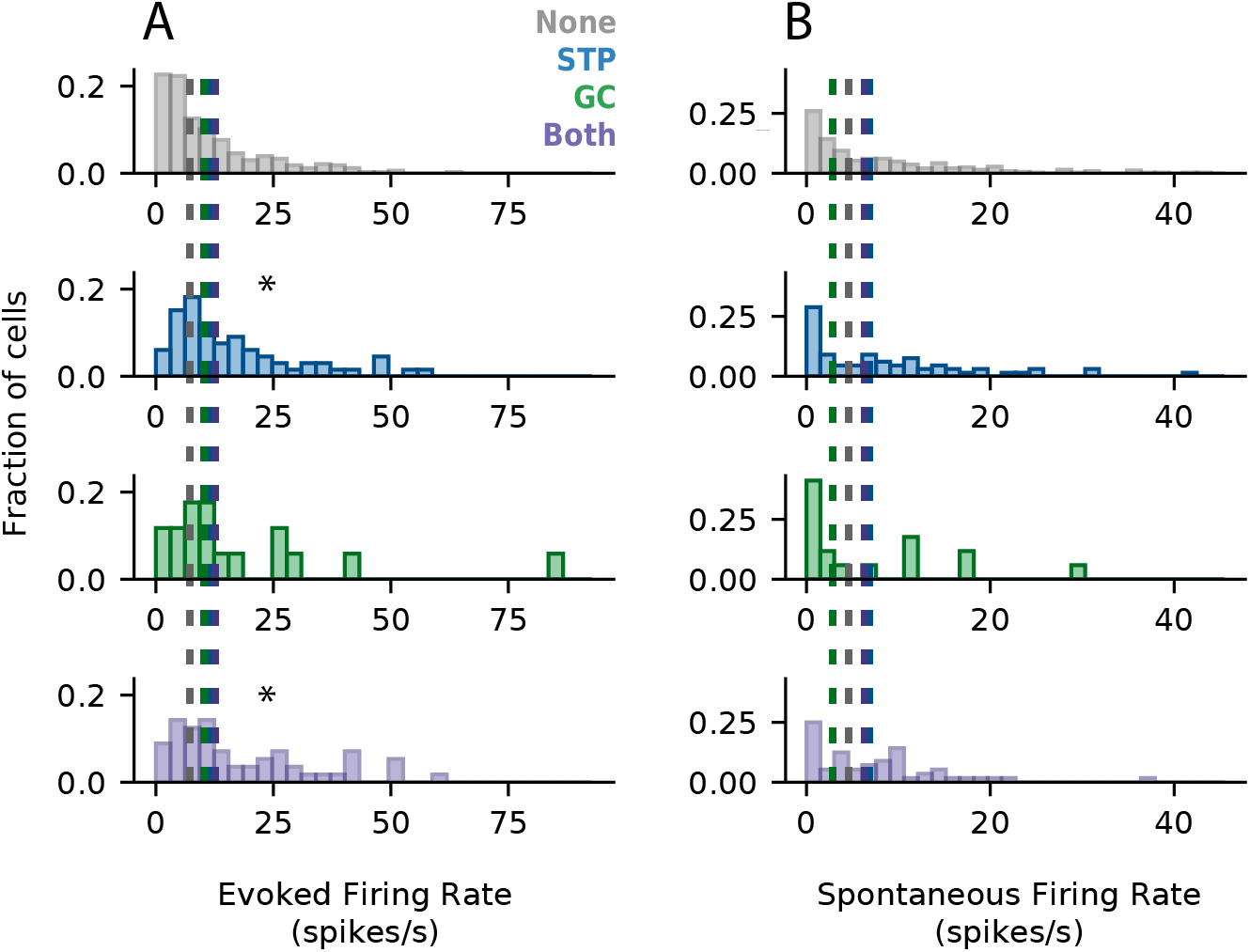
Mean evoked and spontaneous firing rates grouped by nonlinear model performance. (A) Histograms of mean evoked firing rate for four mututally exclusive groups of neurons: no significant improvement in prediction accuracy for the nonlinear models (gray, top), significant improvement for the STP model relative to the LN model (blue, middle-top), significant improvement for the GC model relative to the LN model (green, middle-top), or significant improvement for both the STP and GC models or the GC+STP model (purple, bottom). Median evoked firing rate was significantly higher for the “STP” (11.7 spikes/s) and “Both” (12.7 spikes/s) groups than for the “None” group (7.31 spikes/s, Mann-Whitney U test, *p* = 1.68 × 10^−3^ and *p* = 2.67 × 10^−3^, respectively, adjusted for multiple comparisons). All other comparisons were not statistically significant (*p*^∗^ > 0.05). (B) Histograms of spontaneous firing rate for each model, plotted as in a. None of the groups was significantly different from the others (*p*^∗^ > 0.05).

**Figure 9:**
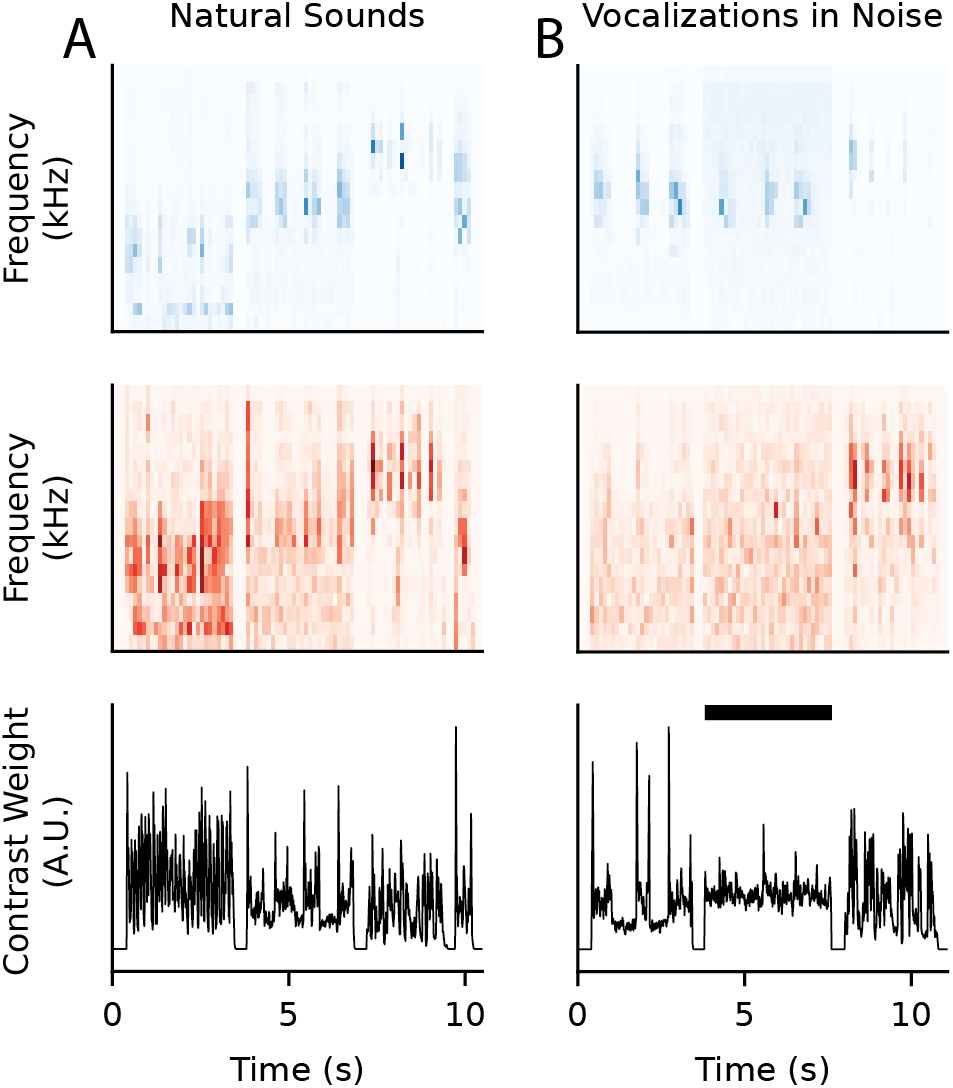
Comparison of natural sound and clean/noisy vocalization properties. (A) Top to bottom: stimulus spectrogram (blue), contrast (red), and frequency-summed contrast (black) for a sequence of three natural sound samples. (B) Same as in A, but for vocalizations. The vocalization set contained interleaved trials of ferret vocalizations with and without additive noise. Black bar indicates segment with noise added.

**Figure 10:**
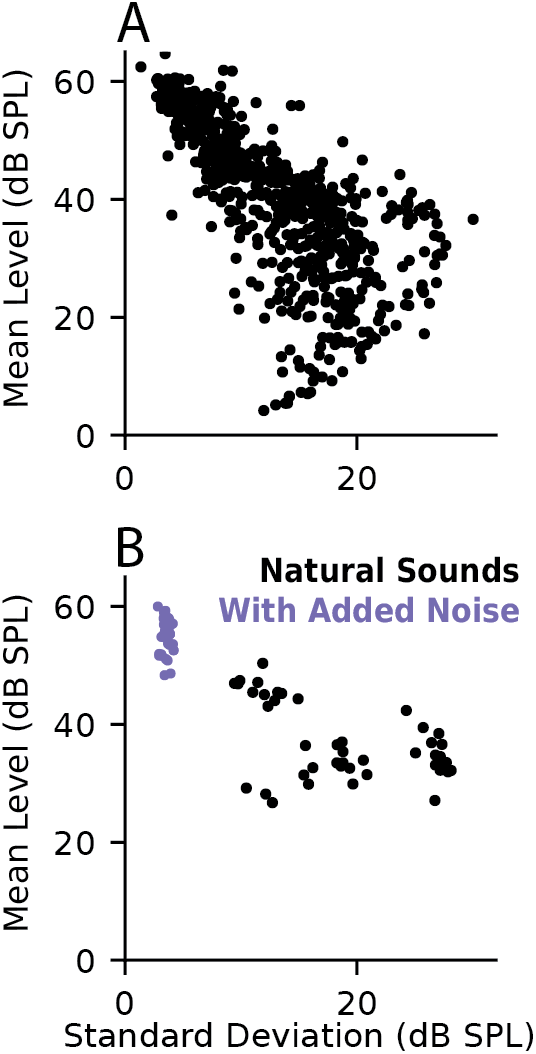
Comparison of contrast properties between stimulus sets. (A) Scatter plot of standard deviation and mean level (dB SPL) for each natural sound spectrogram. The distribution indicates a smooth variation in contrast level. (B) Comparison of standard deviation and mean for clean/noisy vocalizations There is a clear grouping of noisy (low-) and clean (high-contrast) stimuli.

We compared model performance and equivalence, using the same approach as for the natural sound data above (Fig. 11). For the noisy vocalization data, we again found that both the STP and GC models performed significantly better than the LN model and that the combined model performed significantly better than either the STP or GC model individually (Wilcoxon signed-rank tests, two-sided, *p* = 0.0058, *p* = 1.70 × 10^−8^, *p* = 3.69 × 10^−9^, and *p* = 4.27 × 10^−5^, respectively ^*ff,gg,hh,ii*^). However, unlike for the natural sound data, performance of the STP and GC models themselves was not significantly different (Wilcoxon signed-rank test, two-sided, *p* = 0.11 ^*jj*^). This difference indicates a relative increase in the performance of the GC model when applied to noisy vocalizations. This effect is consistent with the hypothesis that gain control plays a bigger role in shaping neural responses for stimuli with large fluctuations between between high- and low-contrast.

**Figure 11:**
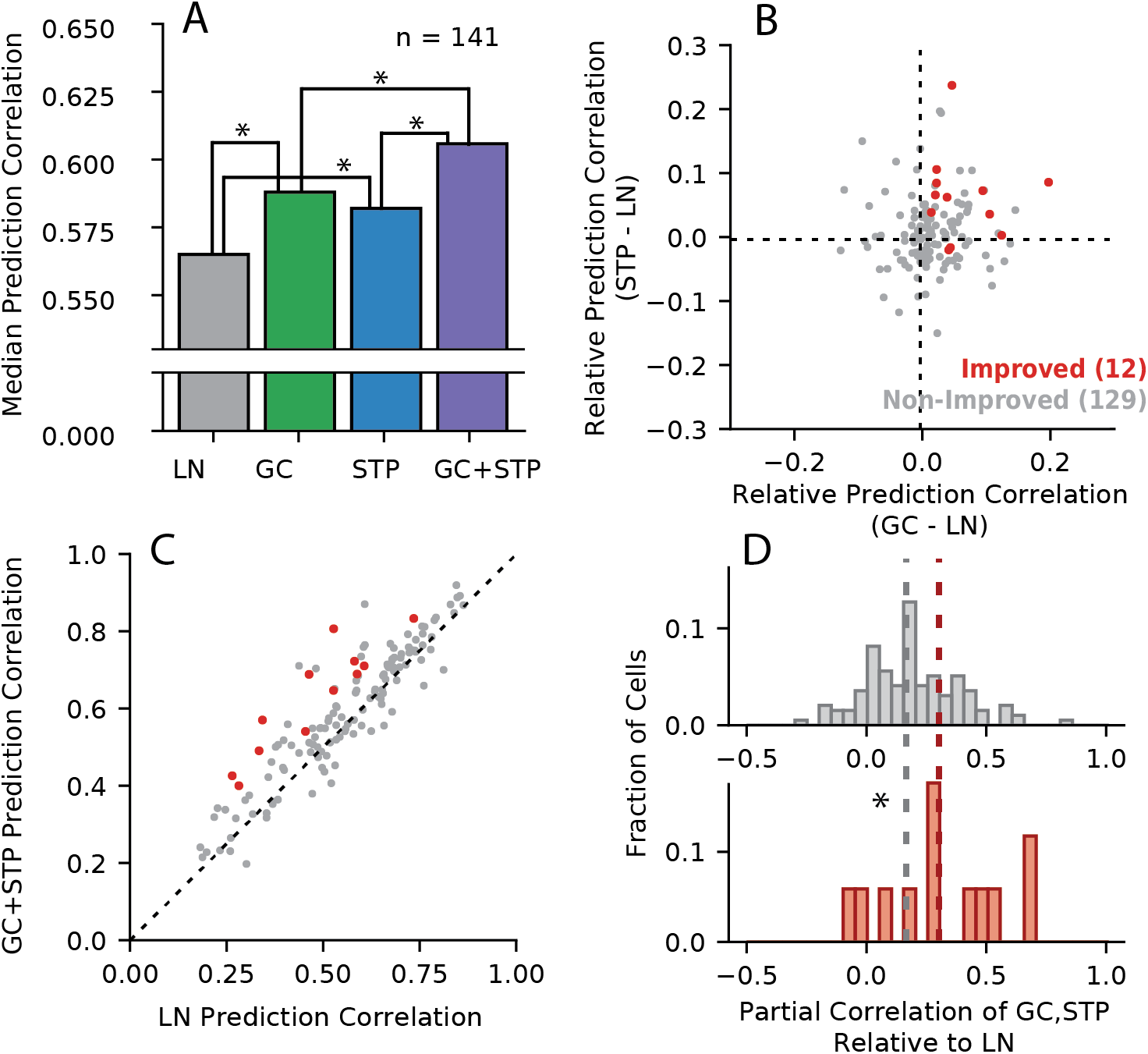
Comparison of model performance for data including clean and noisy vocalizations. (A) Median prediction correlation (*n* = 141) for each model. Statistical significance of median differences was determined via two-sided Wilcoxon signed-rank tests (* *p* < 0.05). Unlike the natural sound data (Fig. 2), performance was not significantly different between the GC and STP models. (B) Change in prediction correlation for the GC (horizontal axis) and STP (vertical axis) models relative to the LN model for each neuron (*r* = 0.18, *p* = 3.45 × 10^−2^). (C) Prediction correlations for the LN model compared to the combined model for each neuron, grouped by whether the combined model showed a significant improvement (red, *n* = 12, *p* < 0.05, permutation test) or not (gray, *n* = 129). (D) Histogram of equivalence for non-improved (top, gray) and improved neurons (bottom, red). Median equivalence for improved cells (0.30) was significantly greater than for non-improved cells (0.18, Mann-Whitney U test, *p* = 0.047).

The equivalence analysis also produced similar results as for the natural sound data. For the noisy vocalizations, the change in prediction correlation was only weakly correlated for the STP and GC models (*r* = 0.18, *p* = 3.5 10^−^2 ^*kk*^). Moreover, the partial correlation between STP and GC predicted responses was modest for improved cells (median 0.30), but significantly higher than the median for non-improved cells (median 0.18, Mann-Whitney U test, two-sided, *p* = 0.047 ^*ll*^). However, the small number of significantly improved cells in this dataset (*n* = 12/141) made drawing definitive conclusions difficult. This smaller set of improved cells likely reflects the fact that the amount of estimation data was smaller for the vocalizations than for the natural sounds.

## Discussion

We found that encoding models incorporating either gain control (GC) or short-term synaptic plasticity (STP) explained complementary aspects of natural sound coding by neurons in primary auditory cortex (A1). Although we observed some degree of equivalence between models, the overlap is modest relative to what would be expected if both models explained the same functional properties (Fig. 4). Instead, a novel model that incorporated both STP and GC mechanisms showed improved performance over either separate model (Fig. 2). It is well-established that the LN model fails to account for important contextual effects (Theunissen et al. 2000; Machens et al. 2004). This work supports the idea that both forward adaptation, mediated by a mechanism such as STP, and feedback inhibition, as might be mediated by GC, both play a role in these contextual processes.

### Assessing equivalence of encoding models

A major goal of this study was to establish a framework for systematically comparing the functional equivalence of complex encoding models. The STP and GC models served as a useful case study for analysis of equivalence: both models account for adaptation following sustained stimulation, and it is not immediately obvious whether they account for distinct contextual effects. By comparing the performance of these alternative models on the same natural sound dataset (Thorson et al. 2015), we were able to determine that they in fact account for distinct functional properties.

Many more encoding models have been developed previously, each representing a separate hypothesis about nonlinear dynamics in auditory encoding (Schinkel-Bielefeld et al. 2012; Harper et al. 2016; Kozlov and Gentner 2016; William et al. 2016; Willmore et al. 2016). In some cases, the equivalence of encoding models can be established analytically Williamson et al. 2015. However, with the development of convolutional neural networks and related machine learning models, future models are likely to only become more complex and difficult to characterize Kell et al. 2018; Keshishian et al. 2019. Alternative models cannot be compared easily because of the many experimental differences between studies in which they are developed. Performance depends not only on the model architecture itself, but also on numerous details of the fitting algorithm and priors that themselves are optimized to the specific dataset of used in a study Thorson et al. 2015. This leaves an important question unanswered: if some number of these models all improve predictive power, does each model’s improvement represent unique progress in understanding the function of auditory neurons or is there overlap in the benefits that each model provides? The equivalence analysis described in this study provides the necessary tools to begin answering this question.

### Increased impact of gain control for stimuli in acoustic noise

The greater relative performance of the GC model for the dataset including noisy vocalizations indicates a behaviorally-relevant regime in which gain control contributes significantly to neural coding in noisy acoustic conditions. The fact that clean and noisy stimuli naturally cluster into high- and low-contrast groups, respectively, may explain the GC model’s increased impact. As a result of this division, a combination of clean and noisy stimuli provides a naturalistic replication of the switches between high- and low-contrast contexts that was used with RC-DRC stimuli in other studies (Rabinowitz et al. 2012; Lohse et al. 2020). In comparison, these binary switches between contrast regimes were absent from the dataset containing natural sounds without added noise, which instead smoothly spanned the contrast space.

In contrast to the variable GC model performance, the consistent performance of the STP model across both data sets suggests that short-term plasticity operates across a wider range of stimuli and is relevant to sound encoding even in the absence of acoustic noise. Previous studies have shown that some nonlinear computations are required in addition to the LN model to account for noise-invariant neural responses in auditory cortex (Moore et al. 2013; Rabinowitz et al. 2013; Mesgarani et al. 2014). The complementary effects of gain control and short-term plasticity reported here are consistent with the idea that both mechanisms contribute to robust encoding of noisy auditory stimuli.

### Comparison with previous studies of gain control

In order to adapt the GC model to analysis of natural sound data, we made some important changes to the original implementation Rabinowitz et al. 2012. First, whereas Rabinowitz et al. imposed the contrast profiles of their stimuli by design, natural stimuli contain dynamic fluctuations in contrast with no predetermined window for calculating contrast. As a result, we were required to make decisions about parameters governing the contrast metric: the spectral and temporal extent of the window used to calculate contrast and the temporal offset needed to emulate the dynamics with which contrast effects are fed back to the response.

We used a 70ms, spectrally narrowband convolution window, which worked best on average in our initial analysis. However, there was variability in the best window for different cells, and model performance may be further improved if these parameters are optimized on a cell-by-cell basis. For the STP model, optimal plasticity parameters also varied substantially between cells. Variability in contrast integration windows may reflect similar biological differences in the ways that cells adapt to contrast. For example, a recent study reported that the timescale of the impacts of contrast on neural gain varied among neurons both in auditory cortex and in two subcortical regions (Lohse et al. 2020).

A second important difference from the original GC model is that we were not able to differentiate high contrast sounds with high standard deviation from those with an exceptionally low mean level—both cases can result in a large coefficient of variation. In the original study, Rabinowitz et al. were able to fix mean level across stimuli in order to avoid this potential confound (Rabinowitz et al. 2012). Despite these differences, however, our results broadly replicated the original findings.

### Mechanisms mediating effects of sensory context on auditory cortical responses

Previous work has described L6 neurons as a mechanistic source of gain control in the visual system (Olsen et al. 2012). Other adaptive mechanisms like stimulus-specific adaptation (SSA) have also been shown to arise from cortical circuits (Natan et al. 2015). However, in the auditory system gain control has been attributed both to cortical feedback and to feedforward adaptation arising in subcortical regions (Lohse et al. 2020). In this case, short-term plasticity could contribute to the feedforward processes (Rabinowitz et al. 2011).

In the current study, the GC model was formulated as a feedback mechanism (Rabinowitz et al. 2012), while the STP model described a feedforward mechanism (David and Shamma 2013). Although we did not directly measure circuit properties, we found that a model combining both mechanisms provided the most accurate predictions overall. Thus our results are consistent with the hypothesis that both contribute to context-related processing in A1. Future experiments involving direct manipulations of synaptic plasticity and/or inhibitory feedback mechanisms can provide explicit insight into the mechanisms underlying these functions.

